# An innovative approach to using an intensive field course to build scientific and professional skills

**DOI:** 10.1101/2021.12.06.471369

**Authors:** Adrienne B. Nicotra, Sonya R. Geange, Nur H. A. Bahar, Hannah Carle, Alexandra Catling, Andres Garcia, Rosalie J. Harris, Megan L. Head, Marvin Jin, Michael R. Whitehead, Hannah Zurcher, Elizabeth A. Beckmann

## Abstract

This paper reports on the design and evaluation of Field Studies in Functional Ecology (FSFE), a two-week intensive residential field course that enables students to master core content in functional ecology alongside skills that facilitate their transition from ‘student’ to ‘scientist’. This paper provides an overview of the course structure, showing how the constituent elements have been designed and refined over successive iterations of the course. We detail how FSFE students: 1. Work closely with discipline specialists to develop a small group project that tests an hypothesis to answer a genuine scientific question in the field; 2. Learn critical skills of data management and communication; and 3. Analyse, interpret and present their results in the format of a scientific symposium. This process is repeated in an iterative ‘cognitive apprenticeship’ model, supported by a series of workshops that name and explicitly instruct the students in ‘hard’ and ‘soft’ skills critical relevant for research and other careers. FSFE students develop a coherent and nuanced understanding of how to approach and execute ecological studies. The sophisticated knowledge and ecological research skills that they develop during the course is demonstrated through high quality presentations and peer-reviewed publications in an open-access, student-led journal. We outline our course structure and evaluate its efficacy to show how this novel combination of field course elements allows students to gain maximum value from their educational journey, and to develop cognitive, affective and reflective tools to help apply their skills as scientists.

## Introduction

Since the 1990s the logistical, resourcing and equity challenges of residential ecology field courses have seen them become increasingly rare in university teaching (Boyle et al., 2007; Cotgreave, 1996). Yet, for university students in ecology, well-structured hands-on activities uniquely build practical research skills (Jackson, 2016) while providing experiences of the excitement and frustration of hypothesis-driven research, data collection and analysis, and collaboration (Abrams et al., 2018; Beckmann et al., 2015; Estavillo et al., 2014; Pedaste et al., 2015). Indeed, field courses are associated with higher self-efficacy gains, higher college graduation rates, higher retention in the ecology and evolutionary biology major, and higher Grade Point Averages at graduation compared to lecture-based courses (Beltran et al., 2020, Scott et al., 2012). The skills attained in field courses also translate to increased graduate employability (Peacock & Bacon, 2018; Mauchline et al., 2013). Studying in the field also helps students understand that nature is incredibly complex, integrated and interdependent, and requires inter-disciplinary thinking (Durrant & Hartmann, 2014; Geange et al., 2021; Behrendt & Franklin, 2014). To maximise the value of learning and the return on investment, therefore, a best practice field course needs to be cost-effective and efficient and provide multiple benefits for both students and teaching staff that extend beyond new discipline knowledge to broader career-enhancing skills.

In this paper, we describe our experiences and evaluations of several years of the Field Studies in Functional Ecology (FSFE) course. During two weeks in the field, coached by experts and peers and supported by appropriately scheduled skills workshops, our students iteratively design and implement customised research projects or ‘field problems’. Students work in small groups to identify their own research questions, design a research protocol, collect and analyse data, and present their findings and interpretations to the group and external stakeholders. Uniquely, student groups develop ‘rapid prototypes’ of a project before swapping it with a new group for refinement and expansion, and after the course can publish their work in an open-access, student-edited journal, closing the research loop through first-hand exposure to scientific publishing.

We designed FSFE to maximise both the value of learning and the return on investment by enabling students to master core content in functional ecology alongside broader employability skills. Boyer (1990), in his seminal work on scholarship, argued that knowledge is acquired through research, synthesis, practice, and teaching. These are all foundational principles in FSFE’s design, not only in the activities provided *for* our students, but also those provided *with* and *by* them, in line with principles of peer learning (O’Donnell & King 1999) and ‘students as [research and teaching] partners’ (Cook-Sather, Bovill & Felten, 2014).Taking hands-on experiences into the field can further add to students’ learning by breaking down the artificial barriers between disciplines (Durrant & Hartmann, 2014; Geange et al., 2021; Behrendt & Franklin, 2014). By providing our students the opportunity to iteratively model the scientific process, while explicitly developing both ‘soft’ and ‘hard’ scientific skills, we provide a unique educational experience that yields professional development as well as rich content delivery.

We aim our course at early-undergraduate students and seek to position our students as active ‘researchers’, as well as students, which allows us to model and shift social identities from ‘students’ to ‘scientists’ (Dennett, 1989). FSFE thus provides a unique educational experience that leads students through an intensive, structured reflective process enabling them to explore their own insights as researchers and peers, yielding richness in both professional development and content delivery. Moreover, inspired by the Organisation for Field Studies field courses (https://tropicalstudies.org/), FSFE has a proven flexibility to work across diverse ecological and environmental biology disciplines and ecosystems. With a broad base of contributing experts and specialists, we have run FSFE in alpine and tropical ecosystems in Australia and in tropical systems in Singapore and Malaysia. Each iteration of FSFE covers the same theoretical principles and scientific concepts but is tailored to location-specific contexts in terms of ecological drivers and locally relevant aspects of protected area management, conservation, and climate change.

### The course structure

The FSFE curriculum has sound pedagogical underpinnings, including achievable learning outcomes aligned with authentic assessment tasks (Biggs & Tang, 2011; **Fig 1)**. The course’s theoretical base lies in cognitive apprenticeship (Brown, Collins & Duguid, 1989): through modelling, coaching, scaffolding, articulation, reflection and exploration (Collins, Brown & Newman, 1987; Enkenberg, 2001; Dennen, 2004), students are ‘apprenticed’ into authentic scientific research practices by the teaching team who explicitly model their expert knowledge and skills in the context of specific learning activities and social collaboration as researchers. We also apply rapid prototyping, whereby scaled-down processes allow faster design, development, evaluation and improvement cycles (Dow & Klemmer, 2011; Garrard et al., 2017).

**Figure 1:**
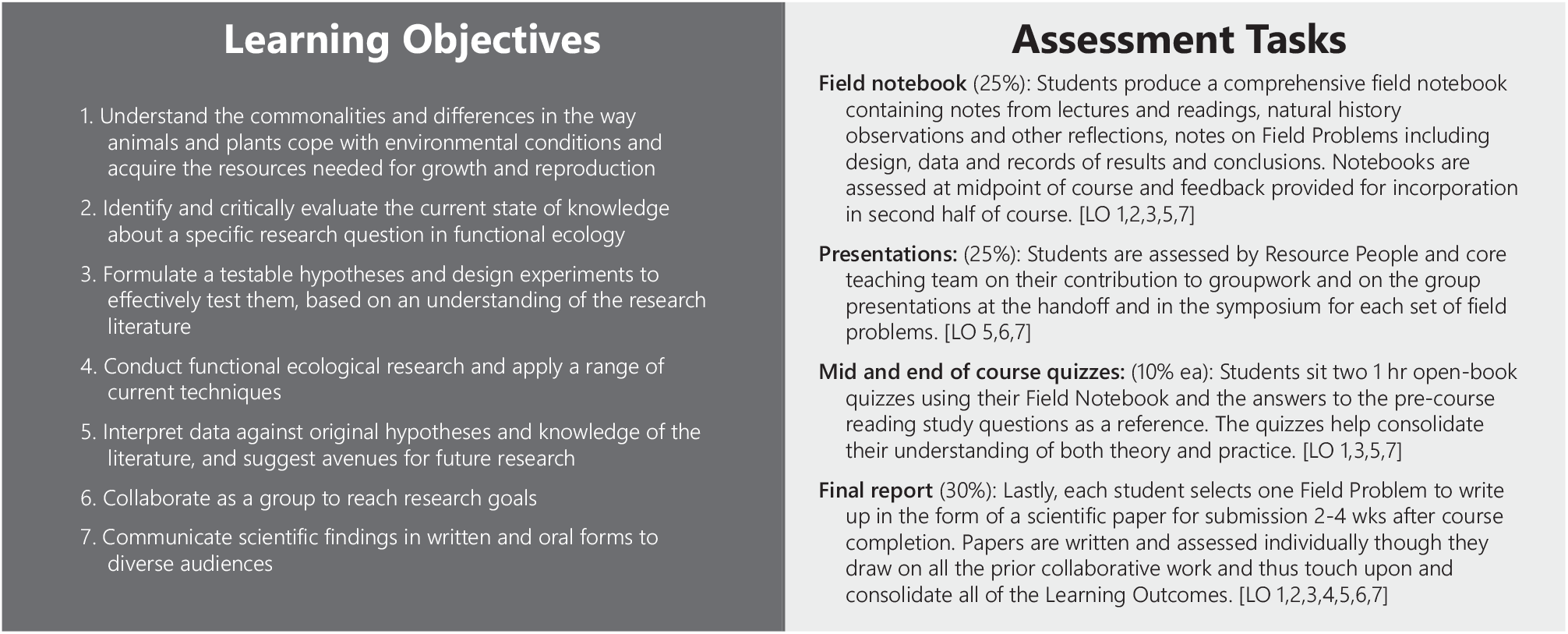
Students are presented with the Learning Outcomes of our course from the outset and Assessment Tasks are aligned to enforce these outcomes. The majority of these are completed during the intensive field component with well-timed feedback so students can reflect on their work and maximise the value of our iterative model.

With active learning a core focus of FSFE, we deliver just 4 lectures that reinforce relevant theory and 10 workshops that present key skills and concepts (**Fig 2a**). These learning activities are all carefully scheduled to meet students’ immediate needs as they develop their projects, acquire data, and then interpret and present their findings (**Fig 1b** and below). Students communicate and refine hypotheses and findings, culminating in a final symposium to which relevant local stakeholders are invited. After the field trip, each student writes a report in the format of a scientific paper, taking time to delve deeper into the literature and cement their learning. Where rigour and quality are sufficient, students are invited to submit their papers in our open-access, student-led journal, where papers are peer reviewed before publication.

**Figure 2a).**
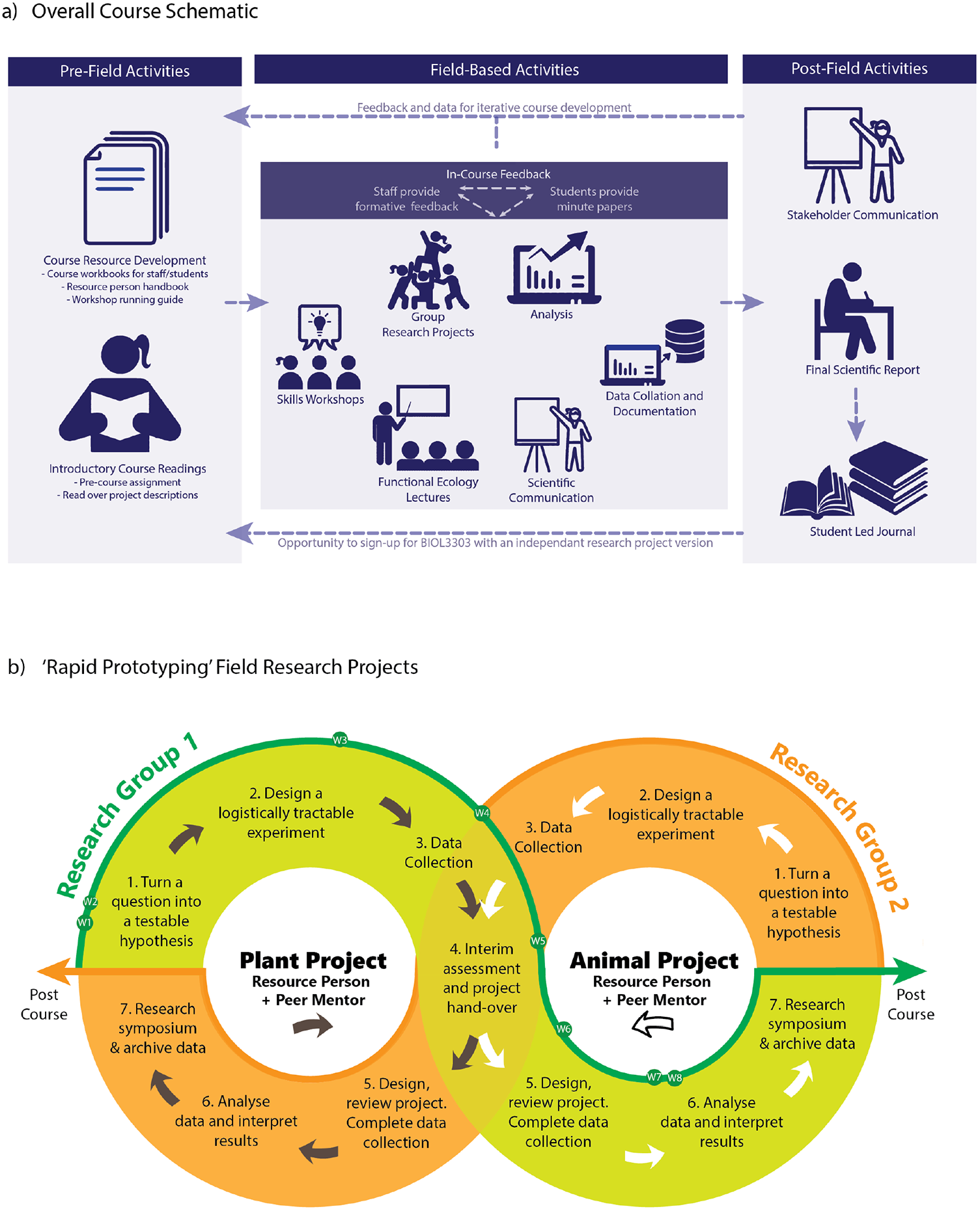
A schematic of overall course structure before, during and after the course. Background content is delivered before the course, the field component aligns skills workshops with phases of the students’ research project, and writing follows the field intensive. b) Illustration of how the students initiate and transfer their projects in each week of the course, showing where each phase of the scientific process applies.

From the outset we designed a companion Advanced version of the course to accommodate a small number of later year students (∼1:3 in comparison to the 2^nd^ year version), including students who had completed the Intermediate-level version. Since 2016, therefore, FSFE has been delivered simultaneously for 2^nd^ and 3^rd^ year undergraduates at Intermediate and Advanced levels respectively. Advanced level students carry out independent research projects developed in consultation with a specialist, and participate in progressive, skill development through parallel Advanced Workshops. Importantly, Advanced students are trained to be peer mentors (**Fig 2**, Workshop 10) for Intermediate student groups, which provides an enriched learning experience for both (Dolan & Johnson, 2009). The detailed course descriptions that follow are based on the Intermediate version of the course.

#### Teaching Team and Specialists

Crucial to the success of the FSFE model are the teaching staff. These comprise two groups—the core teaching and technical team responsible for curriculum, workshop delivery, student pastoral care, planning and logistics; and the transient group of specialists, who we refer to as Resource People.

The core of FSFE is a series of miniature research projects developed by students from their initial exploration of the field environment and supported by the core teaching team and the specialists from diverse ecological disciplines. These specialists assist, coach, model, and advise but do not determine the direction of the research. As our focus is on ecology fundamentals, only a few of our specialists need detailed knowledge of local areas and species. Importantly, the specialists are integrated into the learning as people: as well as their knowledge and teaching, our specialists share their individual perspectives, career and life experiences, contributing to our teaching focus on social and professional identity. This innately human and social perspective helps counter the psychological challenges faced by students as they encounter new concepts, environments and group-dynamics when working in the field, especially in more remote settings (Cotton & Cotton, 2009). Like Goodenough et al. (2015), we have observed that excitement and novelty enhance learning outcomes when students are very well supported.

Overall, some 41 staff have participated, with 6 contributing to three or more iterations, and the same senior academic (course convenor) leading all 7 iterations to date. We recruit the specialists from diverse disciplines, balancing the relative expertise in animals and plants. To enable more researchers to experience the benefits of field-based teaching (Geange et al., 2021), we actively recruit early career researchers into both the core teaching team and as specialists, including Honours or PhD candidates. We embrace high turnover of the specialists as a strength. Past FSFE students are especially welcome.

Most teaching staff have come from our home institution—the Australian National University, Research School of Biology—but we have been privileged to welcome local experts in Far North Queensland, Singapore and Malaysia. Although almost all had some previous university teaching experience, few had previously taught on field courses. All staff are therefore given professional training in field teaching before the course, and constructive support and feedback during the course. Structured evaluations for all staff, in addition to the student evaluations, ensure the core teaching team can act on suggestions from these successive cohorts of specialists, and peer-to-peer mentoring often continues well beyond the course duration.

#### Preparing for the field trip

A month or two before departure, students attend a course induction and Q&A session. They then answer a set of study questions based on pre-course readings that focus their attention on key ecological principles along with course-specific knowledge. Submission of the written responses is a prerequisite for course attendance, though the answers are not graded. For the students, this exercise also provides them with a reference resource during the course, which can be used in the two open-book quizzes.

In preparation for each course, the teaching team and specialists consider study species well in advance, focussing on those we can reliably, legally and ethically investigate in high enough numbers to yield effective sample sizes, given seasonal and weather constraints. To date, projects have focussed on plants, insects, reptiles and birds, all with requisite scientific licenses. On site, we highlight unique or rare flora and fauna in their ecological contexts, and supplement the research projects with appropriate local highlights (e.g., spotlighting for nocturnal arboreal mammal, talks from local land managers, hikes exploring different habitats).

Before the course each of the specialist contributors and most or all members of the core teaching team prepares a “Field Problem Abstract” that poses a question in animal or plant functional ecology—one that interests the specialist and is a genuine open scientific question. Designing projects that can yield a novel discovery and be completed in four days is obviously a challenge. The experienced teaching staff work with the specialists to ensure projects are achievable. Publications in our student-led journal provide examples of what has worked in the past. Students are provided with the compiled Field Problem Abstract Book and an accompanying set of project specific background readings before the course begins. They are encouraged to explore the abstracts but not expected to do any Field Problem specific readings before departure.

#### Field research projects developed on site

Throughout the course the teaching team pay special attention not only to the curriculum but also the students’ mental, social and physical wellbeing. For example, we build the students’ sense of belonging and psychological safety in the first few days by having only the core teaching staff present, before subsequently welcoming specialists to join the group as we move into the Field Problems component of the course. As an ice-breaker, and to ground students’ understanding of multi-disciplinarity and complementary teamwork from the start, we begin with an exercise in which students sort themselves in a line that represents a continuum of their relative interest in plants and animals, in molecular versus landscape perspectives, and their relative confidence with statistics. The exercise of physically mingling amongst the group and learning how their position varies along different axes is an excellent way to meet one another. The teaching team then use the outcomes to allocate students to research project groups that maximise diversity of existing skills and interests.

On day one, guided and mentored by the core teaching team, the students investigate the local ecosystem and begin posing ecological questions and developing test-able hypotheses based on their observations (**Fig 2b, Fig 3**, Workshop 1). On day 2 the students meet the specialists and learn of their group/Field problem allocation (each member of the core teaching team also runs a Field Problem). In this first stage of the process (**Fig 2b**, Research Project 1), each student group works intensively with the relevant specialist to shape a question and hypothesis, and then to design an experiment to test that hypothesis (**Fig 2b**, Research Project 1 Steps 1 to 3).

**Figure 3:**
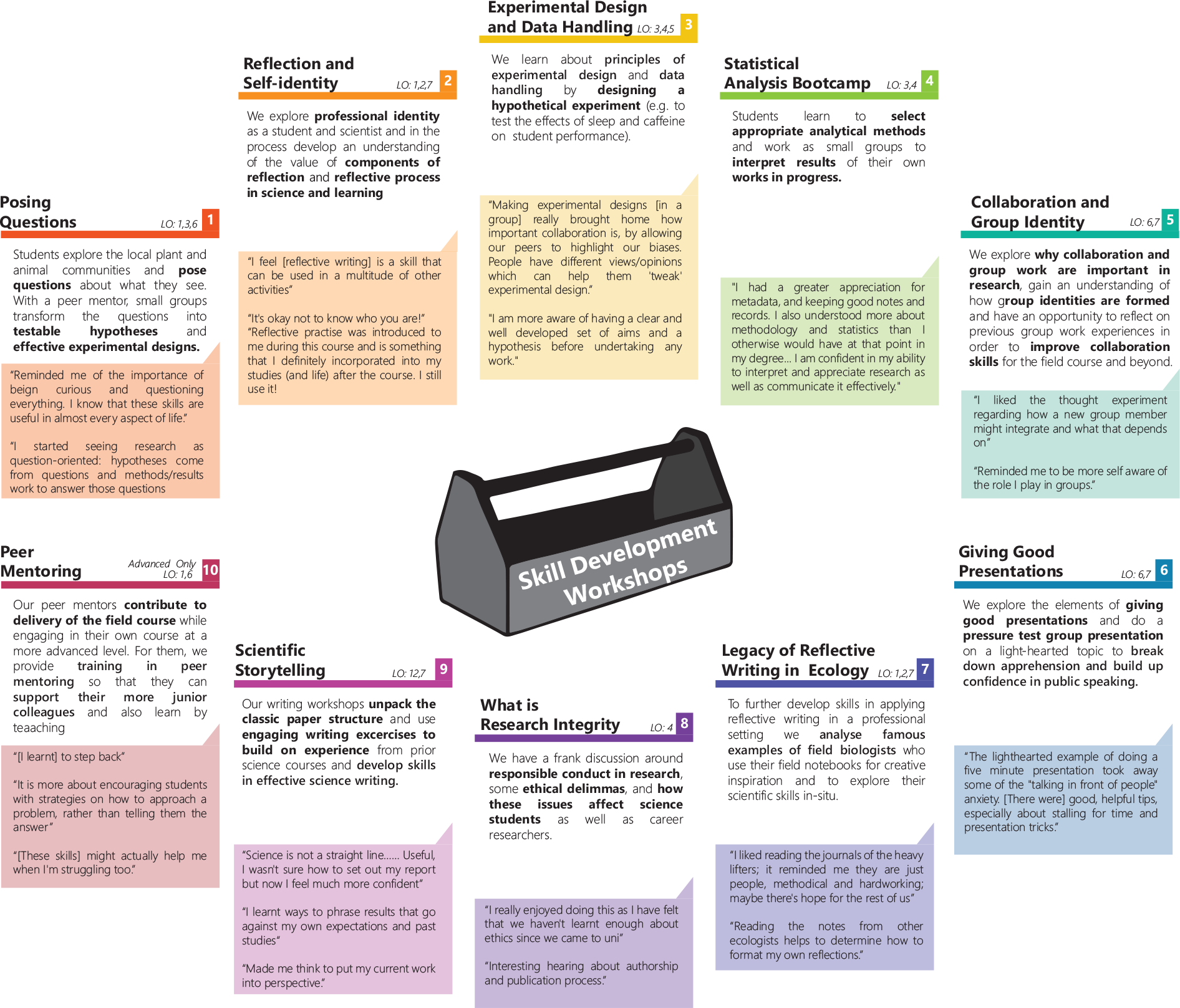
A series of workshops, timed to provide skills at key points in the research process, supports the learning objectives. Quotes drawn from Minute paper evaluations conducted at the end of the workshops demonstrate what the students learn from each

Crucially, the teaching team supports students to frame questions that consider fundamental concepts in functional ecology, can be effectively executed in the field, and generate data that can be analysed at an appropriate statistical level, whatever our location. Students are directed to methods resources (e.g., <u>Prometheus Protocols</u>), but must consider the realities of returns, risks and trade-offs when developing their methods. Approaches range from the simple (e.g., counts of chosen species or measurements of morphological and physical properties) to more advanced (e.g., physiological assays, such as estimating metabolic rates or biochemical constituents). They learn the relative merits of more sophisticated equipment (e.g., leaf gas exchange systems, animal metabolic systems) that generates data at a finer scale but can be difficult to transport and operate in field settings. They discover when a larger sample size might be obtainable using simple, highly reliable equipment (e.g., rulers, binoculars, scales). In so doing, we have enabled students to learn cutting edge techniques and use high-tech equipment in the field. Each student group then initiates their project and conducts 1.5 days of research, before handing the project to a new group (**Fig 1b**, Research Project 1, Step 4).

The handover is one of most unusual and important elements of our course design. At the halfway point of each project, the students swap projects with another group. This handover involves Each group articulating their project’s objectives and hypothesis, and the rationale behind their experiment. Each group also hands over a detailed methods document, a dataset complete with meta-data, and a dot-point summary of the results to date, along with any useful resources (e.g., relevant journal papers or analytical tools). Specialists support the handover process and ensure the students’ research practices meet modern expectations of data archiving and openness (e.g., the FAIR principles described by Wilkinson et al., 2016). Having such a comprehensive hand-over process requires all students to reflect on what they have done and accomplished and tests the data-handling and communication skills of both ‘senders’ and ‘receivers. As students repeat the process during the next research cycle, students can learn from their prior experience how to better facilitate data sharing and learn how to adapt to new collaborations.

Following the handover, each receiving group then decides how to progress the received project: anything from continuing the project unchanged or taking a new approach. As all students are now more familiar with the scientific research process after having begun their own project, the second-round groups tend to be more effective and focused. After another 1.5-2 days of research, students analyse and interpret their data (**Fig 1b**, Research Project 1, steps 5 and 6). Students then present a ∼10-minute conference-style talk on the entire project, including the initial group’s input (**Fig 1b**, Research Project 1, step 7). Lastly, the student archives their data, including meta-data, detailed methods and photos, which ensures that all these resources are available for the write-up phase, and teaches a fundamental principle of modern science.

This whole cycle, from project development to handover and project completion, is repeated for different field problems in the second week of the course (**Fig 1b**, Research Project 2, steps 1 to 7). As the students’ progress through the rapid prototyping cycle (**Fig 1b**), they are continuously prompted to reflect on and develop their skills in collaborative research, including project design and execution, data analysis and interpretation, as well as the oral and written presentation of results.

While this model may appear complex, in practice it flows smoothly, as an elegant example of cognitive apprenticeship strategies in practice. The short-term iterative nature of the rapid prototyping encourages quick design, development, and execution. This maintains a high level of engagement and novelty, encourages students to focus on the fundamentals of scientific research practice, and alleviates pressure on students to obtain conclusive results. Supported by workshops on reflective practice and reflective journals as assessed tasks (see below), the process also explicitly invokes reflective evaluation, consolidation and improvement cycles (Finlay, 2008; Hubbs & Brand, 2005; Kimber et al., 2008; Kolb, 1984; Timpani, 2005).

On the final day of the field course, we revisit all the projects that were conducted, and students are asked to reframe one of their projects in the format of rapid-fire presentation (3 minutes) aimed at a broad stakeholder audience. Relevant local stakeholders (e.g., land managers, conservation practitioners, tourism operators) are invited, to hear what the students have learned and to provide them with feedback on their ability to communicate their work to a lay or stakeholder audience. This final presentation is voluntary and not assessed, but almost invariably all the students engage with the exercise in some way and find the presentations a fitting way to celebrate their accomplishments.

#### In-field workshops provide an explicit focus on skills development

Skills Workshops are a key element of the teaching in FSFE (**Fig 3**). In addition to the usual scientific process skills, we explicitly name and build ‘soft’ skills—interpersonal strengths, communication, emotional intelligence, reasoning, and problem-solving skills—that are highly sought by employers in any field (Graduate Careers Australia, 2016; Laker & Powell, 2011; Mauchline et al., 2013; Peatland et al., 2019). By making this part of the course explicit, we find that student demonstrate have a better appreciation of why we include the workshops and a greater sense of ownership of their learning (Stokes, Mangier & Weaver, 2011). The workshops are mandatory and one hour long, with most held in the late-afternoon before students have free time and dinner or occasionally in the evening, as after-dinner fare. The workshops are structured around clear objectives (Beckmann et al., 2017), interactive engagement, and summary handouts for students. Regularly updated facilitator handbooks, slide presentations, optional handouts and relevant equipment enables facilitators to deliver to the same high standards even if they are new to the teaching team. Students provide immediate post-workshop feedback via 1-minute papers (**Fig 2**), which has enabled continuous refinement of content and delivery.

Six Skills Workshops centre on helping students unpack the scientific process (**Fig 3**). An initial ‘Posing Questions’ workshop (W1) familiarises students with the local flora and fauna and helps them convert observations and curious questions into testable hypotheses. When each student group has framed its hypothesis, we consider experimental design and data handling (W3). Focused thinking about applied statistics occurs near the end of their first Field Problem (W4). As students prepare their first of several oral presentations, W6 pairs public speaking skills with light-hearted improvisation activities. Towards the end of the course, science writing skills are explored (W9). Finally, we dedicate a session to considering research integrity, moving beyond the normal lectures admonishing plagiarism and instead introducing students to the complexity of scientific authorship, ethical considerations around research and data handling, and the codes of practice that inform professional research (W8). For most students this first exposure to ethical practice beyond the issue of plagiarism has proved a revelation.

Four additional Skills Workshops focus on building a researcher identity and developing skills in collaboration and reflective practice; these are key course goals related to the cognitive apprenticeship model (**Fig 3**). These innovative workshops build on affective learning as a strong component of field courses (Boyle et al., 2007; Beckmann et al., 2017). At the start of the course, we explore the concepts of personal reflective practice and the ‘social identity approach’ (Haslam, 2004) in relation to behaviour within and between teams of collaborating researchers (W2). Through regular entries in the Field Notebooks students reflect on their participation and the course as part of an experiential learning cycle (Kolb, 1984; Moon, 1999) with a view to gaining insight into themselves as learners and scientists. After the first project students are also able to reflect on their own and their team’s challenges and strengths as collaborators, so we extend our discussion by exploring how students and research scientists, and ecologists in particular, build self-identity and complementary teamwork skills (W5). A focus on reflection as part of research practice comes next (W7). Knowing that students might conflate reflective journals with simple diaries, we explore the field journals of notable naturalists and ecologists to demonstrate how reflection on field observations and notes have contributed historically to the development of ecological theories. This helps students see their Field Notebook assessment task more holistically.

#### After the trip

By the end of the course, each student has participated in four different projects (two on animals and two on plants), one of which they select to write up as their final paper due 2-4 weeks after return. The write-up draws on the methods, data and presentation materials that were put in our archive during the course. This both models a key element of contemporary science and gives the students maximum flexibility in writing up their final paper (Gallagher et al., 2021; Parker et al., 2016). The final papers are written individually, in the style of a research paper, and follow the format of our student-led journal.

Throughout FSFE, we emphasise researchers’ responsibility to communicate and ideally publish their findings to maximise potential impact (US National Research Council, 2003). Student-led undergraduate journals are relatively rare, especially in science, yet are known to provide particularly powerful learning, especially if peer review experiences are included (Guilford, 2001; Uigín, Higgins & McHale, 2015). From the first iteration of FSFE, we inaugurated an open access journal ‘*Field Studies in Ecology*’ (**Fig 4;** see supplement **S1** for more detail). Students who achieved a ‘Distinction’ (∼>70% or a B) for their final report can choose to submit a manuscript for peer review. The expectations for these junior authors are high: they need to show a substantive understanding of relevant discipline knowledge, critical thinking, and data analysis and synthesis, and scientific writing, alongside thoughtful responses to feedback from the expert peer reviewers. The journal’s editors are also FSFE students, selected for each volume through expressions of interest along with academic performance. Mentored by academics and professional editors, these student editors take on significant responsibilities in the peer review and publication process, including managing all student authors and academic peer reviewers selected from the current and former specialists, colleagues in our Research School, and where relevant external researchers.

**Figure 4.**
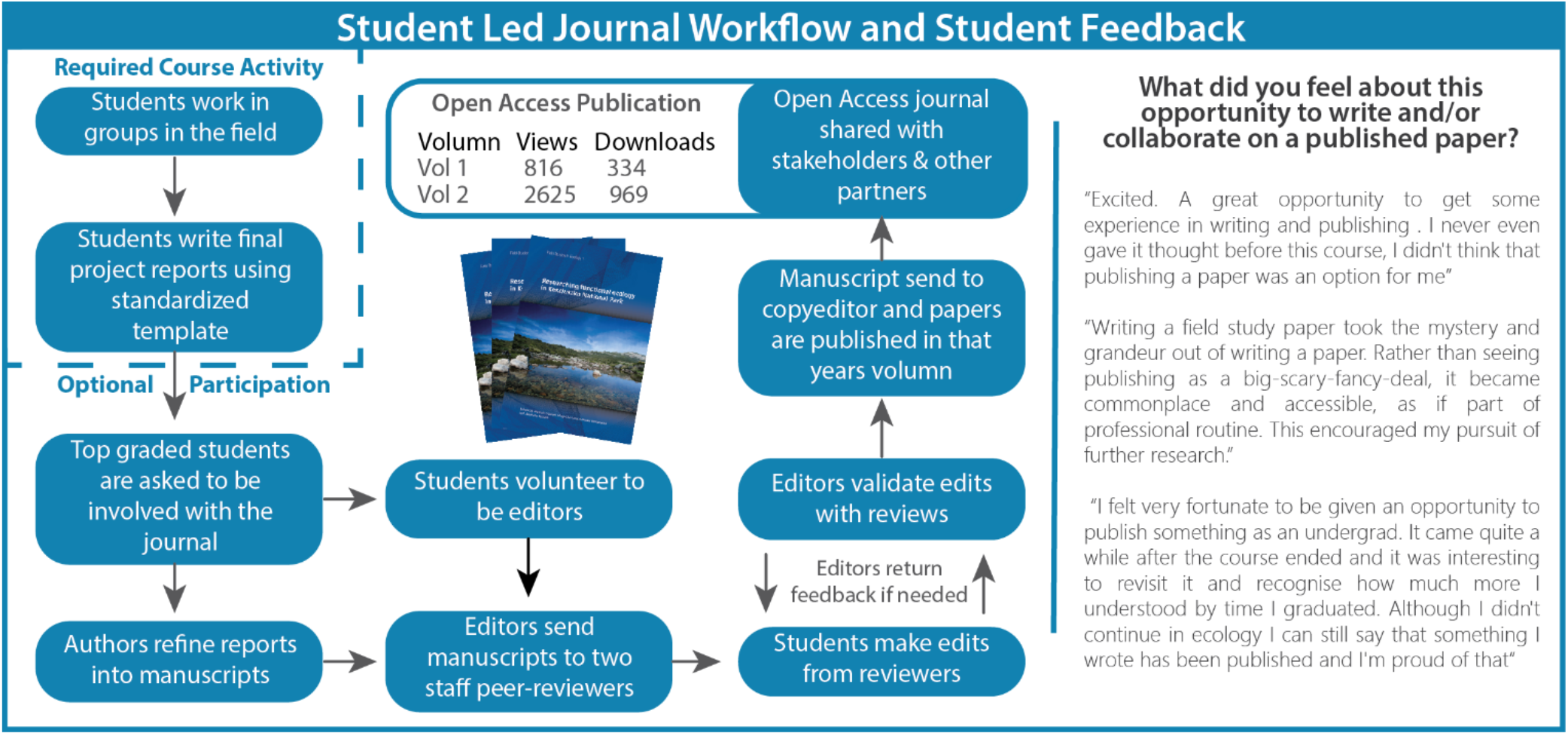
Field Studies in Ecology is an open-access, student-edited, peer-reviewed journal that makes student research accessible for subsequent students, stakeholders, and the broader community while also giving the students genuine experience of the process of scientific writing, review and publication.

Four volumes of ‘*Field Studies in Ecology*’ have been published to date. Volumes 1 and 2 respectively cover the 2015 and 2016 courses at Kosciuszko National Park (Zurcher et al., 2017; Hazell-Pickering, Slatyer & Nicotra, 2019). Volume 3 includes research from 2017 at the Daintree Rainforest (Cape Tribulation, Queensland) and 2018 at the Bukit Timah Nature Reserve in Singapore (Harris et al., 2021). Volume 4 is in preparation. The journal makes the FSFE student data available to stakeholders—government agencies, land managers, industry professionals. Analytics show all volumes are viewed and downloaded regularly (**Fig 4**).

### Assessing the effectiveness of the FSFE model

A continuous improvement cycle (Temponi, 2005) has been a feature of all facets of the course since the first iteration. In 2020, we sought evidence about the longer-term impacts of FSFE in three ways. First, we surveyed all students and staff who had participated in the course using an online survey that included Likert-scale questions on FSFE’s impact on the students’/staff areas of interest, knowledge and skills, and open-ended questions on perceptions of the impact of FSFE overall. Second, we collated a random sample of 85 paired reflective writing entries from Field Notebooks, written in the middle and at the end of the course, and assessed the relative development of reflective practice. Entries were assessed across the four attributes associated with effective reflective writing—descriptive detail, emotive engagement, critical reflection, and meta-reflection – using the assessment rubric that we provide to the students (Moon, 1999; Kember et al., 2008). Third, we compared the academic outcomes of our students using a paired student design where students are compared to a student from the same degree and who achieved the same grade in the pre-requisite first year course who did not complete FSFE. The results are summarised in **Fig 5** and as answers to the four questions below (further detail on methods and analyses is available in **S2)**.

**Figure 5:**
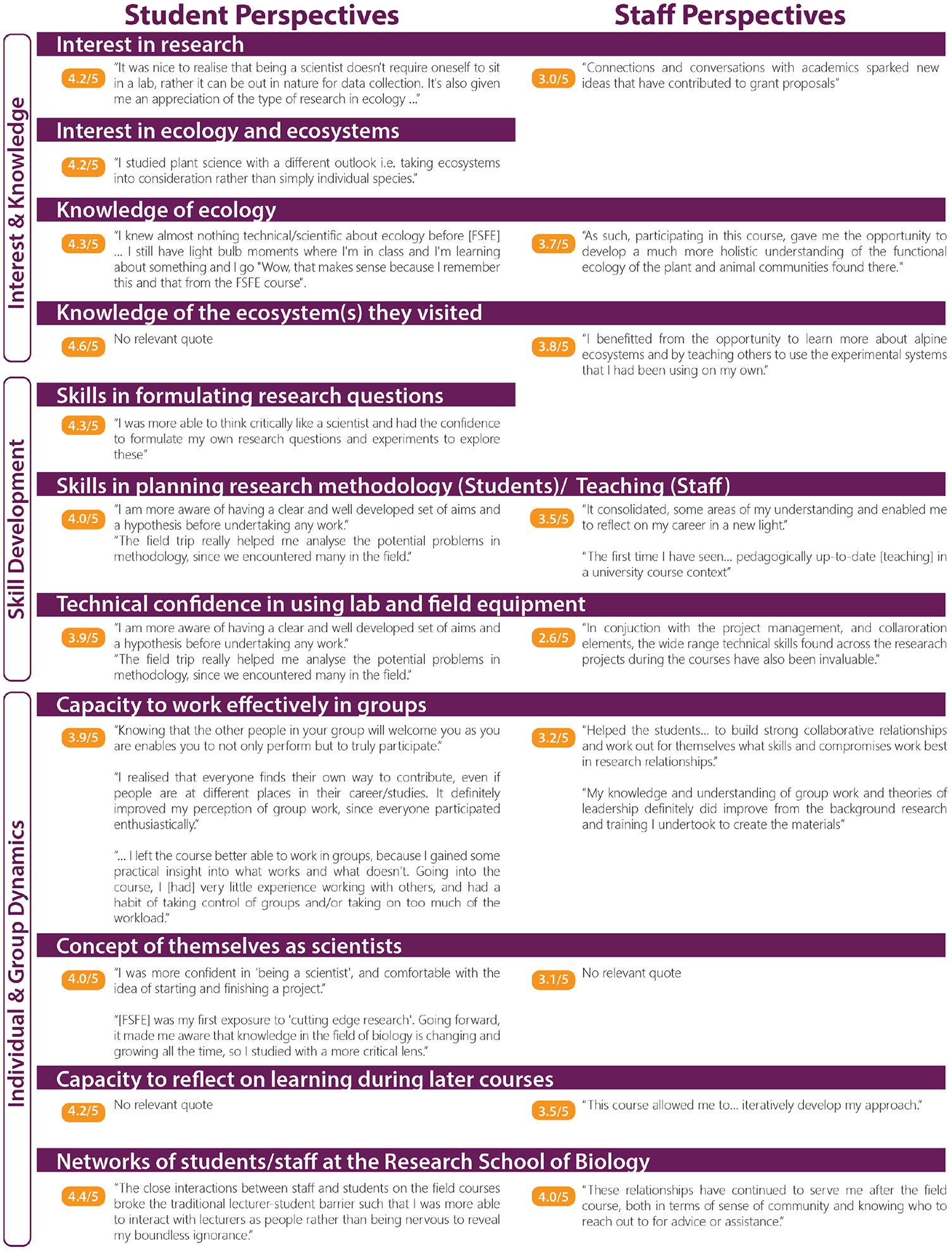
Responses of students (left column) and staff (right) to a retrospective survey of perceptions of the impact of FSFE. The survey questions used a Likert-scale ranging from 1 (not at all) to 5 (very much). Mean ratings above 3/5 were considered

#### How has FSFE influenced students’ study and career paths?

Although FSFE delivers an unusually high standard of research skills and opportunities, it was not designed to only serve students seeking research careers in ecology. Our quantitative analysis showed that students who took FSFE (either as a second year, third year or both) were more likely to complete their degree (noncompletion of FSFE students = 3.5% versus noncompletion of non-FSFE students = 17.6%, see S1). However, we are unable to determine whether this is a consequence of taking FSFE or due to an intrinsic property of the students who take FSFE (i.e., students that choose to take a course like FSFE are more likely to finish their degree). In the student survey, respondents reported substantial impacts of their FSFE experience on their subsequent studies and interests (**Fig 5**). For example, FSFE had motivated some to take more biology and ecology courses in their science degree than they had originally planned, and many acknowledged FSFE as having motivated them to pursue research careers. Indeed, most (70%) of the 25 student survey respondents who had graduated since taking FSFE had continued into research (Honours, Masters programs, Doctor of Philosophy) or further study (Doctor of Medicine degrees), and others were working in related fields—science communication, government science policy, or non-university ecology research. Most (76%) reported that FSFE had given them confidence to identify themselves as scientists, regardless of their future career directions.

Regardless of their subsequent study and career paths, almost all the respondents to the student survey reported that they had approached learning and community-building differently after FSFE. The novel settings, the student-led approach, and the opportunities for high quality interactions with staff stood out for many as specific features of the course. Many reported that FSFE provided their first experience of ‘real science’—authentic exploration and discovery. Although our quantitative comparison of their outcomes in terms of GPA did not reflect a statistically significant impact given the small sample size (see **S1**), students reported that the improved confidence and skills they felt they had gained during the course positively impacted their subsequent academic performance.

Students described how being supported through the scientific process in FSFE had enabled them to think more critically, assimilate information with greater ease, and develop specific scientific/academic skills that benefited their future study. In open-ended responses, many described the lasting impact of FSFE: these students felt the course had affected them in ways that still permeated both their personal and academic lives, stimulating memories and reflections years later.

#### Do FSFE students develop skills in research and scientific thinking?

Almost all student survey respondents reported notable increases not only in their *knowledge* of ecology and the ecosystems they visited but also in their *interest* in ecology and ecosystems, and in research overall. Some two thirds reported notable increases in their ongoing skills in formulating research questions and planning research methodology after FSFE, while a similar proportion noted increased technical confidence in using lab and field equipment. The knowledge-sharing in FSFE often occurs ‘just in time’ i.e., in relation to genuine curiosity and a ‘need to know’. Our students reported they could assimilate a much greater amount of complex content in this learning context compared to the equivalent taught in packaged lectures in a campus-based delivery model.

#### Do FSFE students develop skills in reflective practice?

As educators we wanted to know whether FSFE was improving students’ capacity for reflective practice. As we had provided both training and written mid-course feedback on each students’ individual reflections, we hypothesised a comparison would reveal that students’ reflective proficiency and competencies would improve over the course. Students scored consistently high on descriptive detail in their reflective writing from the beginning of each course, which is expected given the students were motivated and this is the most basic attribute of reflective writing (Moon, 1999). By contrast, by the end of the course scores had significantly increased for increasingly sophisticated and more effective reflective practice including emotive engagement with their experiences (P=0.01), ability to critically reflect, evaluate and analyse (P<0.001), and ability to reflect on the value of reflection (meta-reflection, P<0.001). Overall, and especially in students who attended both Intermediate and Advanced iterations, the journals demonstrated a clear shift from ‘reflection *on* action’ to ‘reflection *in* action’ which Schon (1983) considered the core of ‘professional artistry’, and Findlay (2008) described as indicating an expert who acts “both intuitively and creatively [as they] revise, modify and refine their expertise”. This analysis of the reflective journal data was supported by the student survey findings: for example, most respondents (76%) reported a greater capacity to reflect on their own learning in their subsequent university courses after completing FSFE.

#### Do FSFE students develop teamwork skills?

Another key focus of FSFE is collaborative teamwork. In each iteration, we have observed tangible improvements in teamwork. For example, the teaching team regularly observe that individuals become better at drawing on the diverse skills and capabilities of their peers as well as of the specialists, and at sharing responsibilities, outcomes, and discoveries. The reflective culture adds to this by facilitating a greater awareness and tolerance of their own and their peers’ limitations.

In the student survey, most respondents (83%) reported that FSFE had increased their capacity to work effectively in groups. Responses to open-ended questions showed that these FSFE participants felt that being supported through the scientific process had enabled them to think more critically, assimilate new information with greater ease, and develop specific scientific/academic skills that benefited their future work and study. Most respondents (83%) also reported that FSFE had initiated a notable growth in their networks of peers and staff.

#### FSFE staff evaluation of the teaching model

Lastly, we assessed the feedback from 21 staff who responded to the 2020 survey for their insights on the teaching model for students and for their own professional development. The staff reported that FSFE students benefited from the course’s applied, immersive nature: needing to be realistic in data collection and project design/management had given the students a ‘real’ experience of practicing science. Irrespective of previous teaching experience, all staff respondents also reported that participating in FSFE had improved their teaching skills, especially their capacity for reflecting on their teaching practice. Staff also commonly reported that teaching in FSFE had increased their technical confidence and broader knowledge of ecology, enlarged their professional networks and research collaborations, increased the number of students seeking supervision in research degrees, and had provided unique opportunities to give and receive mentorship.

### Closing remarks

Our evaluation of five years of FSFE has shown a great array of positive outcomes including reports of increased self-efficacy, learning gains, confidence, collaboration skills, research interest and more among our students. Pairing ecological content and skills development workshops with our rapid prototyping of research projects under ‘apprenticeship’ to specialists has proven highly effective. Explicitly weaving concepts of social identity and reflective practice into the course, and a commitment to teaching complex ‘soft’ skills like collaboration, reflective practice and teamwork has had clear benefits. The measured shift from reflection ‘on’ action to reflection ‘in’ action indicates individuals more capable of recognising, and engaging with, the diverse skills and capabilities of any cohort. Further, by continuously evaluating and fine tuning our model, FSFE has become a highly effective and novel teaching tool that delivers major, positive impacts on student academic and professional development.

FSFE is a vehicle for learning, teaching and practising authentic contemporary science, including data sharing, peer review and open access publishing. The FSFE field course model provides an outstanding vehicle for research-led education that finds and nurtures talent, actively engages with stakeholders beyond the university, and fosters collaborative and reflective practice that is preparing students to address the pressing real-world challenges. Imbued with the intentional perspective that we all share a scientist identity. This course further facilitates a gentle but enduring identity transition from ‘we are a group of students and academics’ to ‘we are all scientists creating knowledge together’.

We hope that our exploration of how the FSFE course functions as in its evolving form provides inspiration for development of other field courses. Through field courses our students gain authentic research experience and come to appreciate both what the skillset of a scientist is and what the value of those skills is in a diverse array of successful careers. Maintaining field courses in an undergraduate curriculum can be challenging due to the logistical constraints and costs, but these courses are so important to developing skills that improve graduate employability that they are crucial to ensuring well-rounded undergraduate experiences (Mauchline et al., 2013).

## Acknowledgements

The authors would like to acknowledge the RSB leadership for supporting the course’s development and especially supporting our inclusion and diversity model by providing student bursaries from 2016. The authors would like to acknowledge all the other staff who have contributed to the design and delivery of FSFE since 2015, and of course all the students who have participated. We would particularly like to thank the indefatigable Wes Keys for proving that all this was logistically possible: no worries. Our thanks also to Panit Thamsongsana and the RSB Biology Teaching and Learning Centre for ongoing support and encouragement.

The data collection from students and staff reported in this paper was authorised under The Australian National University Human Research Ethics Protocol 2015/553.

Our course framework is easily modified and has already been adopted and adapted, by other universities. Supporting materials for this course (including Field Problem Abstract Books and workshop materials) are available on request.

## Author Contributions

All authors made substantial contributions to conception and design, acquisition of data or analyses. All authors contributed to writing the paper and approved final submission.

## Data accessibility statement

Data will be made publicly available on FigShare as allowable under human ethics constraints.

## Figure Legends

Figure 1. A schematic of a) the overall course structure pre, during and post-intensive and b) the rapid prototyping approach of the field problems.

Figure 2. Students are presented with the Learning Outcomes of our course from the outset and Assessment Tasks are aligned to enforce these outcomes. A series of workshops, timed to provide skills at key points in their research process, supports the learning objectives. Quotes drawn from Minute Paper evaluations (Stead, 2005) conducted at the end of the workshops demonstrate what the students learn from each.

Figure 3. : Responses of students (left column) and staff (right) to a retrospective survey of perceptions of the impact of FSFE. The survey questions used a Likert-scale ranging from 1 (not at all) to 5 (very much). Mean ratings above 3/5 were considered to indicate an increase in the relevant sphere. Open ended comments from the survey are included to illustrate impacts. Blank spaces on the staff side indicate that a question was not asked of the staff.

## Literature cited

Abrams, D., Masser, B., Houston, D., & McKimmie, B. (2018). A social identity model for education. In: Argote, L., & Levine, J. M. (Eds.). The Oxford Handbook of Group and Organizational Learning. Oxford University Press. https://doi.org/10.1093/oxfordhb/9780190263362.013.1

Beckmann, E. A., Estavillo, G. M., Mathesius, U., Djordjevic, M. A., & Nicotra, A. B. (2015). The plant detectives: Innovative undergraduate teaching to inspire the next generation of plant biologists. Frontiers in Plant Science, 6, 729. https://doi.org/10.3389/fpls.2015.00729

Beckmann, E. A., Weber, X., Whitehead, M., & Nicotra, A. (2017). Research-based learning: Designing the course behind the research. pp. 141–151. In: Zurcher, H., Ming-Dao, C., Whitehead, M., & Nicotra, A. (Eds.). Researching Functional Ecology in Kosciuszko National Park. Field Studies in Ecology. Volume 1. ANU eVIEW. https://doi.org/10.22459/RFEKNP.11.2017.14

Behrendt, M., & Franklin T. (2014). A review of research on school field trips and their value in education. International Journal of Environmental and Science Education, 9(3), 235–245. https://doi.org/10.12973/ijese.2014.213a

Beltran, R. S., Marnocha, E., Race, A., Croll, D. A., Dayton, G. H., & Zavaleta, E. S. (2020). Field courses narrow demographic achievement gaps in ecology and evolutionary biology. Ecology and Evolution, 10(12), 5184–5196. https://doi.org/10.1002/ece3.6300.

Biggs, J., & Tang, C. (2011). Teaching for quality learning at university: What the student does. Society for Research into Higher Education & Open University Press.

Boyer, E. L. (1990). Scholarship reconsidered: Priorities of the professoriate. Princeton University Press.

Boyle, A., Maguire, S., Martin, A., Milsom, C., Nash, R., Rawlinson, S., Turner, A., Wurthmann, S., & Conchie, S. (2007). Fieldwork is good: The student perception and the affective domain. Journal of Geography in Higher Education, 31(2), 299–317, https://doi.org/10.1080/03098260601063628

Brown, J. S., Collins, A., & Duguid, P. (1989). Situated cognition and the culture of learning. Educational Researcher, 18(1), 32–42.

Collins, A., Brown, J. S., & Newman, S. (1989). Cognitive apprenticeship: The teaching of reading, writing and mathematics. In L.B. Resnick (Ed.), Knowing, Learning and Instruction. Essays in Honor of Robert Glaser (pp. 453–494). Lawrence Erlbaum.

Cook-Sather, A., Bovill, C., & Felten, P. (2014). Engaging students as partners in learning and teaching: A guide for faculty. Jossey-Bass (Wiley).

Cotgreave, P. (1996). Fertile fields of study. Times Higher Education, May 17. https://www.timeshighereducation.com/news/fertile-fields-of-study/93654.article

Cotton, D. R. & Cotton, P. (2009). Field biology experiences of undergraduate students: the impact of novelty space. Journal of Biological Education, 43(4), 169–174. https://doi.org/10.1080/00219266.2009.9656178

Dennen, V. P. (2004). Cognitive apprenticeship in educational practice: Research on scaffolding, modeling, mentoring, and coaching as instructional strategies. In D. H. Jonassen (Ed.), Handbook of Research on Educational Communications and Technology. (pp. 813–828). Lawrence Erlbaum.

Dennett, D. C. (1989). The intentional stance. MIT Press.

Dolan, E., & Johnson, D. (2009). Toward a holistic view of undergraduate research experiences: An exploratory study of impact on graduate/postdoctoral mentors. Journal of Science Education and Technology, 18, 487. https://doi.org/10.1007/s10956-009-9165-3

Dow, S. P. & Klemmer, S. R. (2011). The efficacy of prototyping under time constraints. pp. 111–129 In: Plather, H., Meinel, C., Leifer, L. (Eds). Design Thinking: Understand, Improve, Apply. Springer.

Durrant, K. L. & Hartman, T. P. V. (2015) The integrative learning value of field courses. Journal of Biological Education, 49(4), 385–400, https://doi.org/10.1080/00219266.2014.967276

Enkenberg, J. (2001). Instructional design and emerging teaching models in higher education. Computers in Human Behavior, 17(5-6), 495–506.

Estavillo, G. M., Mathesius, U., Djordjevic, M., & Nicotra, A. B. (2014). The plant detective’s manual: A research-led approach for teaching plant science. ANU Press. https://doi.org/10.22459/PDM.11.2014

Finlay, L. (2008). Reflecting on ‘reflective practice’. Practice-based Professional Learning Paper 52. The Open University.

Gallagher, R. V., Falster, D. S., Maitner, B. S. et al. (2021). Open Science principles for accelerating trait-based science across the Tree of Life. Nature Ecology & Evolution, 4, 294–303. https://doi.org/10.1038/s41559-020-1109-6

Garrard, G. E., Rumpff, L., Runge, M. C., & Converse, S. J. (2017). Rapid prototyping for decision structuring: An efficient approach to conservation decision analysis. pp. 46–64. In: Bunnefeld, N., Nicholson, E., & Milner-Gulland, E. J. (Eds.), Decision-Making in Conservation and Natural Resource Management: Models for interdisciplinary approaches. Cambridge University Press.

Geange, S. R., von Oppen, J., Strydom, T., et al. (2021). Next-generation field courses: Integrating Open Science and online learning. Ecology and Evolution, 11, 3577–3587. https://doi.org/10.1002/ece3.7009

Goodenough, A. E., Rolfe, R.N., MacTavish, L. & Hart, A.G. (2015). The role of overseas field courses in student learning in the Biosciences. Bioscience Education. https://doi.org/10.11120/beej.2014.00021

Graduate Careers Australia. (2016). Graduate Outlook 2015. The report of the 2015 Graduate Outlook Survey: Perspectives on graduate recruitment. https://www.graduatecareers.com.au/files/wp-content/uploads/2016/07/graduate-outlook-report-2015-final1.pdf

Guilford, W. H. (2001). Teaching peer review and the process of scientific writing. Advances in Physiology Education, 25(1-4), 167–75. https://doi.org/10.1152/advances.2001.25.3.167.

Harris, R., Nix, S., Head, M. & Posch, B. (Eds). (2021). Field Studies in Ecology. Volume 3. ANU eVIEW. https://studentjournals.anu.edu.au/index.php/fse/issue/view/21

Haslam, S. A. (2004). Psychology in organisations: The social identity approach. London: Sage.

Hazell-Pickering, S., Slatyer, R., & Nicotra A. B. (Eds.) (2019). Researching functional ecology in Kosciuszko National Park. Field Studies in Ecology. Volume 2. ANU eVIEW. https://studentjournals.anu.edu.au/index.php/fse/issue/view/13

Hubbs, D. L., & Brand, C. F. (2005). The paper mirror: Understanding reflective journaling. Journal of Experiential Education, 28(1), 60–71. https://doi.org/10.1177/105382590502800107

Jackson, D. (2016). Re-conceptualising graduate employability: The importance of pre-professional identity. Higher Education Research & Development, 35(5), 925–939.

Kember, D., McKay, J., Sinclair, K., & Wong, F. K. Y. (2008). A four-category scheme for coding and assessing the level of reflection in written work. Assessment & Evaluation in Higher Education, 33(4) 369–379, https://doi.org/10.1080/02602930701293355

Kolb, D. A. (1984). Experiential learning: Experience as the source of learning and development. Volume 1. Prentice-Hall.

Laker, D. R. & Powell, J. L. (2011). The differences between hard and soft skills and their relative impact on training transfer. Human Resource Development Quarterly, 22, 111–122. https://doi.org/10.1002/hrdq.20063

Mauchline, A. L., Peacock, J. & Park, J. R. (2013). The future of bioscience fieldwork in UK higher education. Bioscience Education, 21(1), 7–19. https://doi.org/10.11120/beej.2013.00014

Moon, J. (1999). Reflection in learning and professional development: Theory and practice. Routledge-Falmer.

O’Donnell, A. M., & King, A. (1999). Cognitive perspectives on peer learning. Lawrence Erlbaum.

Parker, T. H., Forstmeier, W., Koricheva, J., Fidler, F., Hadfield, J. D., Chee, Y. E., Kelly, C. D., Gurevitch, J., & Nakagawa, S. (2016). Transparency in ecology and evolution: Real problems, real solutions. Trends in Ecology and Evolution, 31(9), 711–719. https://doi.org/10.1016/j.tree.2016.07.002

Peacock, J. & Bacon, K. L (2018). Enhancing student employability through urban ecology fieldwork. Higher Education Pedagogies, 3(1), 440–450. https://doi.org/10.1080/23752696.2018.1462097

Peasland, E.L., Henri, D.C., Morrell, L.J. & Scott, G.W. (2019). The influence of fieldwork design on student perceptions of skills development during field courses. International Journal of Science Education, 41(17), 2369–2388. https://doi.org/10.1080/09500693.2019.1679906

Pedaste, M., Mäeots, M., Siiman, L. A., de Jong, T., van Riesen, S. A. N., Kamp, E. T., Manoli, C. C., Zacharia, Z. C., & Tsourlidaki, E. (2015). Phases of inquiry-based learning: Definitions and the inquiry cycle. Educational Research Review, 14, 47–61. https://doi.org/10.1016/j.edurev.2015.02.003

Schon, D. A. (1983). The reflective practitioner. Basic Books.

Scott, G. W., Goulder, R., Wheeler, P., Scott, L.J., Tobin, M. L., & Marsham, S. (2012). The value of fieldwork in life and environmental sciences in the context of higher education: A case study in learning about biodiversity. Journal of Science Education and Technology, 21(1), 11–21. https://doi.org/10.1007/s10956-010-9276-x

Stead, D. R. (2005). A review of the one-minute paper. Active Learning in Higher Education, 6(2), 118–131. https://doi.org/10.1177/1469787405054237

Stokes, A., Magnier, K. & Weaver, R. (2011). What is the use of fieldwork? Conceptions of students and staff in geography and geology. Journal of Geography in Higher Education, 35(1), 121–141. https://doi.org/10.1080/03098265.2010.487203

Temponi, C. (2005). Continuous improvement framework: implications for academia. Quality Assurance in Education, 13(1), 17–36. https://doi.org/10.1108/09684880510578632

Uigín, D. N., Higgins, N., & McHale B. (2015). The benefits of student-led, peer-reviewed journals in enhancing students’ engagement with the academy. Research in Education, 93(1), 60–65. https://doi.org/10.7227/RIE.0010

US National Research Council. (2003). Committee on Responsibilities of Authorship in the Biological Sciences. Sharing Publication-Related Data and Materials: Responsibilities of authorship in the life sciences. 2. The purpose of publication and responsibilities for sharing. National Academies Press. https://www.ncbi.nlm.nih.gov/books/NBK97153/

Wilkinson, M. D., Dumontier, M., Aalbersberg, I., et al. (2016). The FAIR Guiding Principles for scientific data management and stewardship. Scientific Data, 3, 160018. https://doi.org/10.1038/sdata.2016.18

Zurcher, H., Ming-Dao, C., Whitehead, M., & Nicotra, A. B. (Eds.). (2017). Researching functional ecology in Kosciuszko National Park. Field Studies in Ecology. Volume 1. ANU eVIEW. https://doi.org/10.22459/RFEKNP.11.2017

