## Supplementary File 1 for "An innovative approach to using an intensive field course to build scientific and professional skills"

### Nicotra et al.
### Supplement 1: The student led journal

##### Box 1: From Student Assignment to Peer-Reviewed Publication: *Field Studies in Ecology*

Our students have the opportunity to publish the research conducted during the FSFE courses in the peer-reviewed, open access journal '*Field Studies in Ecology*' (https://studentjournals.anu.edu.au/index.php/fse). The first two volumes cover the 2015 and 2016 courses respectively at Kosciuszko National Park (Zurcher et al, 2017; Hazell-Pickering et al, 2019) while the third volume also includes research at the Daintree Rainforest Observatory at Cape Tribulation, Queensland, in 2017 and at the Bukit Timah Nature Reserve in Singapore in 2018 (Harris et al, 2021). Volume 4 is in preparation. This body of published work reflects the values of student-led learning and is driven by students as both the primary researchers and journal editors. This continuous engagement—from initiating the project idea in the field through to publication—develops students' first-hand engagement with the peer-review process, an aspect of scientific communication rarely available to undergraduates or even very early career researchers (Guilford, 2001; Uigín, Higgins & McHale, 2015).

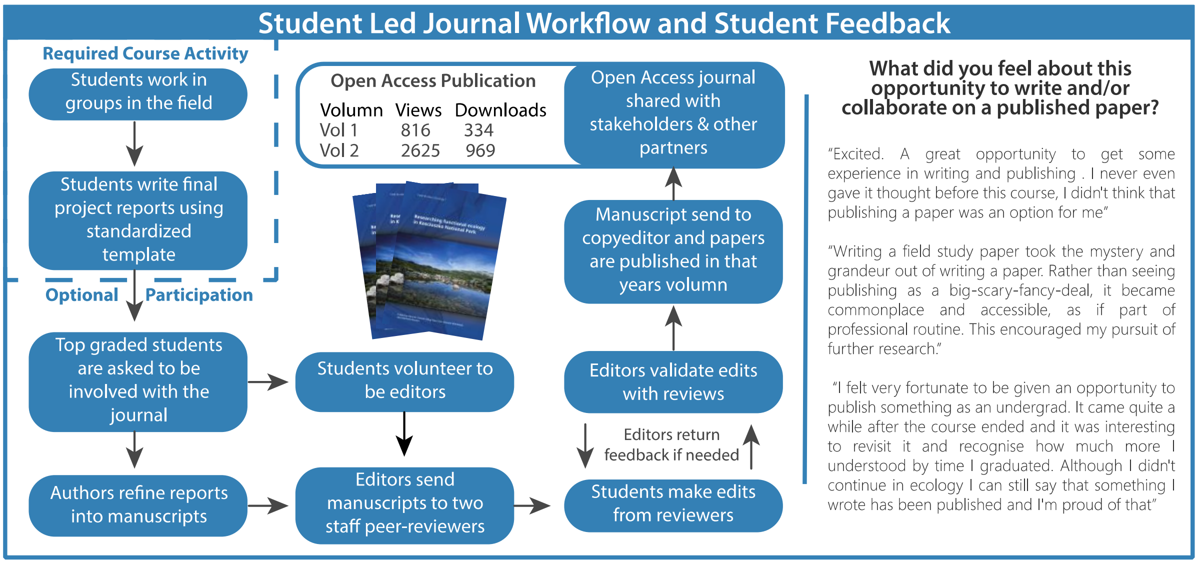


**Student editors:** Applicants for the journal editor roles come from the FSFE cohorts and are subjected to a rigorous selection process. Mentored by academics and professional editors as they develop their skills, the student editors are given a high level of responsibility in handling the whole peer review and publication process. This includes managing all the student authors and selecting and overseeing a large number of peer-reviewers, including many senior academic researchers. The leadership skills that the editors develop are proving an exceptional asset to their career paths, whether they continue in academia or move into other professional areas.

**Submissions:** Course assessment requires students to submit a final report based on one of their field projects. To nurture and reward enthusiasm, effort and quality, students who do this to a good standard (usually graded at 70%, a Distinction of ‘B’, or more) are offered the option of submitting their manuscript for traditional peer-review, rewriting to an even higher standard, and consideration for publication in the journal. We only accept one paper per Field Problem, so if more than one student wants to submit for the same small-group project, we support them to collaborate in joint authorship.

The journal is hosted at ANU on the Open Journal Systems software platform. The system facilitates the submission and review process and looks, acts, and feels like any other journal front end.

**Undergraduate authors:** To achieve publication, the student authors must demonstrate a high level of mastery of relevant discipline knowledge and research capabilities. The peer review feedback naturally strengthens these aspects, as well as bolstering the student authors’ skills in written communication, critical thinking, data analysis and synthesis, all key employability skills. The process also facilitates career thinking about the realities of academia.

**Research engagement and outreach:** In addition to its role as an educational tool, the journal aspires to contribute more widely by making available data and thinking on the ecologies of some vulnerable ecosystems—addressing a specific gap in the publication space. The journal is not only open-access but is actively shared with many stakeholders, including other scientists, government agencies, community groups, land managers and industry professionals. Data analytics show that the journal is viewed and downloaded regularly.

**Student feedback:** Students’ reflections on their experience with the journal have been collected from the course-required reflective journals and the recent participant survey. These data show that the opportunity to publish research as undergraduates was not only unexpected but quite transformative. Students overwhelmingly underestimated their capacity to contribute, and then demonstrated a more than sufficient understanding of requirements to produce work of high quality. In the process students gained a sense of achievement and satisfaction from working as a group and being able to publish a body of work from their cohort. Even when asked some years later, students reported that the exercise developed their identity as legitimate researchers, illustrated the benefits of teamwork, and gave them an inordinate sense of pride.
