## Supplementary File 2 for "An innovative approach to using an intensive field course to build scientific and professional skills"

### Nicotra et al.

### Supplement 2: A detailed description of our evaluation of the efficacy of the FSFE Model

#### Introduction

As already noted, we have maintained a consultative continuous improvement cycle (Temponi, 2005) across all facets of the course since the first iteration. This has included impacts noted in students' reflective journals and minute papers during each iteration giving insight into in-course outcomes, as well as proximate post-course evaluations giving more holistic views.

In their recent large-scale study at University of California Santa Cruz, Beltran et al (2020) found that field courses were associated with higher self-efficacy gains (more so than lecture-based courses), higher college graduation rates, higher retention in the ecology and evolutionary biology major, and higher Grade Point Averages at graduation. While recognising that our much smaller sample size would not necessarily yield statistically strong quantitative analyses, we certainly anecdotally recognised self-efficacy gains in our FSFE students, and therefore embarked upon a retrospective evaluation to provide a more detailed understanding of the impact of the FSFE experience on students' learning outcomes. To better assess the long term impacts of participation in FSFE we combined retrospective surveys, assessment of development in reflective practice during the course, and an analysis of impacts on academic outcomes.

For a long term perspective we assessed the effectiveness of the FSFE approach more objectively via a retrospective survey of students up to 5 years after completing the course. In 2020 (when the course was cancelled because of COVID-19 circumstances) we surveyed all former students, teaching team and resource people who had participated in FSFE since its inauguration. This allowed us to consider especially the longer-term impact of FSFE on students' career trajectories, which is of special interest given the course's focus on both the scientific method and student identity as scientists.

Enabling our students to become more active reflective learners was a key goal of FSFE. As already noted, students were trained in reflective practice and writing during the course (Figure 2, W2 and W7), and their reflections were assessed at the course mid-point and at the end of the course. We therefore analysed whether students improved in their capacity for reflective practice during the course. Firstly, anecdotally, those of us reviewing and assessing students' field notebooks across the two weeks of each course observed clear shifts in the quality and depth of students' reflections, moving towards the higher-level attributes that Moon (1999) associates with effective reflective writing. Moreover, from our conversations on site to the survey responses one to five years after participation, we observed students (especially in those who returned for the advanced course) clearly shift from 'reflection on action' to 'reflection in action' which Schon (1983) considered the core of 'professional artistry', and Findlay (2008) described as indicative of the expert who acts "both intuitively and creatively [as they] revise, modify and refine their expertise".

Finally, though at our university FSFE could account for usually only one, and at most two, of the required 24 courses for graduation with a Bachelor of Science, we conducted a quantitative analysis to see whether participation in FSFE had any measurable effect on academic outcomes. We compared academic results and outcomes across Science of FSFE participants compared with non-participants. We anticipated that the absolute impact on performance measures, e.g. GPA, would be small but that participation may reflect choices that the students made in terms of area of study, whether or not they proceeded to further study, and what their career choices were.

Here we present, as a companion to our Ecology and Evolution Academic Practice article a detailed assessment of our evaluation process and its outcomes.

### Methods

#### Assessment of impact of FSFE on student academic outcomes

As our sample of FSFE students was not random, we compared academic outcomes of the FSFE cohorts relative to their peers using a dataset of paired student contrasts (n=105 pairs). Here, we paired each FSFE student with another student who had not taken the course, but who had achieved the same grade in the same first year prerequisite course and was enrolled in the same degree. We did not account for other potential covariates such as whether students were first generation or non-English speakers for this analysis as we did not have those data.

Using Generalized Linear Mixed Models (GLMM) implemented in R statistics package “lme4” (R Core Team, 2021; Bates et al, 2015), and the paired student design based on anonymised academic transcripts, we assessed whether taking part in FSFE affected the final overall Grade Point Average (GPA) of graduates, whether graduates were more likely to complete their degree and whether they were more likely to continue onto a research study pathway (i.e. enrol in Honours (an undergraduate research year in Australia) or postgraduate study). In each case, we specified GLMM with whether or not a student participated in the FSFE course as a fixed categorical effect, and pair identity as a random effect. For the analysis of final GPA, we used a Gaussian error distribution and power transformed the response variable (GPA) to ensure model residuals were normally distributed. The appropriate power (3.62) to use was determined using the power. Transform function in the R package “car”. For the analysis of the likelihood of degree completion and continuing on a research pathway we specified a binomial error distribution with a logit link.

#### Retrospective student and staff survey

In late 2020 we contacted all students who had participated in at least one of the six FSFE iterations using university email addresses. Noting that these might no longer be accessed by graduates, reducing potential response rates and sample size, we also relied on word of mouth and course-specific Facebook groups. We invited the students to respond to an online survey in which we asked open-ended questions on their perceptions of the impact of FSFE, and Likert-scale questions on whether FSFE had stimulated increases in specific areas of interest, knowledge or skills (See **Fig 5,** and Supplement **S3**).

We also asked contributing staff to respond to an online survey with Likert-scale and open-ended questions that, where appropriate, paralleled those asked in the student survey.

##### Developing skills in reflective practice

To assess development of skills in reflective practice we collated a random sample of 85 mid/end reflective journal entries from across all course iterations (in 2017 a reflective question on the mid and end of course quiz was assessed instead of the journal) and both Intermediate and Advanced students. Four experienced members of the teaching team then scored the students' reflective writing across four attributes—descriptive detail, emotive engagement, critical reflection, and meta-reflection (after Moon, 1999; Kember et al, 2008)—with a grade from 0 (none shown) to 4 (excellent evidence). We analysed the mid/end score pairs with ordinal logistic regression in R-package 'MASS' (Venables and Ripley 2002). As we provided both the cohort training and written mid-course feedback on each students' individual reflections, our hypothesis was that students' reflective proficiency and competencies should show signs of improvement over the course.

#### Results & Discussion

##### Assessment of impact of FSFE on student academic outcomes

We found that completion of FSFE was significantly associated with increased degree completion rates, although number of non-completions (3.5% among FSFE students and 17.6% among those who did not take FSFE, $\chi^2 = 26.251$, $p < 0.001$) was small overall. However, we found no association between completion of FSFE (either the Intermediate or Advanced version) and going on to further postgraduate or Honours study (34.3% of FSFE students relative to 32.4% of non) ($\chi^2 = 0.115$, $p = 0.734$). We also found no effect of participation in an FSFE course on the final overall GPA of graduating students (mean GPA ± SE: FSFE student = 5.830 ± 0.085, non-FSFE students = 5.806 ± 0.098) ($\chi^2 = 0.063$, $p = 0.803$).

##### Retrospective student and staff survey

###### Student responses

The sample of 43 respondents was spread evenly across course iterations (each represented by responses from 8-13 former students) and course level (86% had participated at Intermediate level and 45% at Advanced level, with 30% having taken both). All respondents were enrolled in a Science degree, with 24% also enrolled in a non-Science degree (double degree). When they first attended FSFE, 60% were in the first year of their Science degree, 20% in their second year, 17% in their third year and 2% in their fourth or later year. About 60% of survey respondents had graduated since taking FSFE. Of these, the majority (72%) had continued into research or further study (50% enrolled or planning to enrol in Honours or Masters programs, 20% enrolled or planning to enrol in a Doctor of Philosophy program and 12% enrolled in Doctor of Medicine degrees). Of the other graduates, 24% were working in science communication or government science policy, while 12% were employed in non-university ecology research.

Respondents reported substantial impacts of their FSFE experience on their subsequent studies and interests (**Fig. 5**). The novel settings, the student-led approach, and the opportunities for high quality interactions with staff stood out for many as specific features of the course itself. Many reported that FSFE provided their first experience of 'real science' and that they were sure the improved confidence and skills they gained during the course positively impacted their subsequent academic performance.

One specific set of survey questions asked how much FSFE had stimulated increases in various aspects of relevant interest, knowledge or skills, with potential responses presented on a Likert scale ranging from 1 (not at all) to 5 (very much). Mean ratings above 3 were considered to indicate an increase in the relevant sphere (Figure 3). Almost all respondents reported notable increases not only in their *knowledge* of ecology and the ecosystems they visited but also their *interest* in ecology and ecosystems, and in research overall. Some two thirds reported notable increases in their skills in formulating research questions and planning research methodology post-FSFE, and a similar proportion noted that their technical confidence in using lab and field equipment had increased.

> *... discussing our [possible research] questions was extremely rewarding and engaging as I was both learning and collaborating with others .... I felt different after the group discussion*
> *knowing it was one of my questions that we are going to test tomorrow.* (Extract from reflective journal, 2018)

In addition, most respondents reported that FSFE had increased their capacity to work effectively in groups (83%, mean 4.0/5) and gave them confidence to identify themselves as scientists (76%, mean 4.0/5). Responses to the open-ended questions clearly showed that these FSFE participants felt being supported through the scientific process had enabled them to think more critically, assimilate new information with greater ease, and develop specific scientific/academic skills that benefited their future study.

Most respondents (83%, mean 4.4/5) also reported a notable growth in their networks of peers and staff as a result of their participation in FSFE.

> *It was revolutionary to ... find myself in a group full of people just like me, who'd stay up ... going through the literature and fixing the presentation ... who didn't mind that I'd done the reading, who'd done the reading themselves, who would actually discuss what we'd read.* (Participant in 2015 & 2016, BA/BSc, now working in government policy development)

> *I learnt to accommodate more work styles in order to achieve a shared goal* (2015 & 2016, 2nd year student, graduated BSc, now technical officer in research agency)

> *I was better able to organise myself to work with ... group members after the field trip ... useful on some of the larger projects I had later in my degree, through finding out the best division of labour to each task.* (2017, 1st year student, graduated BSc, planning Masters of Ecology)

> *I found it easier to work in groups after learning more strategies of ways in which groupwork could be assigned and completed. I learnt that everyone has something*

> *unique to offer during groupwork, and that even if you do not have a perfectly cohesive group, that group is not forever.* (2019, 2nd year student, enrolled BSc)

The responses from the survey demonstrated very clearly that these former students—some of whom had taken the course five years earlier—felt that they had approached learning and community-building quite differently after FSFE. Most respondents (76%) reported that post-FSFE they experienced a greater capacity to reflect on their own learning in their subsequent university courses. Specifically, they described how being supported through the scientific process enabled them to think more critically, assimilate information with greater ease, and develop specific scientific/academic skills that benefited their future study. Some reported specifically that taking FSFE had motivated a shift in their study trajectory toward taking more biology and ecology courses, and many reported that FSFE had increased their motivation to pursue further research, which was evidenced in their post-FSFE study/career paths. In open-ended responses, many described the lasting impact of the course: they felt participating in FSFE had affected them in ways that still permeated both their personal and academic lives, stimulating memories and reflections even years later.

##### Staff responses

We surveyed staff to compare their assessment of the impact and efficacy of involvement in FSFE for students, as well as for their own professional development. About half of the 21 staff who responded were in early- to mid-career stages, and a few had themselves initially been students in FSFE.

When asked to reflect on what they considered the value of the course for students, staff respondents were unanimous in their view that FSFE students benefited from the course's applied, immersive nature. There was a particular recognition that the practice-based need for students to be realistic in their approach to data collection and project design/management while on site had given the students a 'real' experience of practicing science.

All 21 staff respondents reported having some kind of university teaching experience before participating in FSFE, with most (85%) having tutoring/demonstrating experience in on-campus courses and 30% having been course convenors. These respondents' professional interest in teaching was evident: almost half (45%) had been involved in peer discussions about teaching, some had completed teaching development programs (35%), and some (25%) had received professional recognition in teaching, such as national or institutional teaching awards and/or fellowships of the Higher Education Academy (25%).

However, just 15% reported having already been highly experienced in supporting residential field courses—which we defined as having taught in four or more field courses before FSFE. Notably, 60% reported having had no experience in teaching or supporting students at residential field courses before their participation in FSFE. While these findings are unsurprising given the ongoing global cost-cutting shift away from field-based teaching that started in the 1990s (Boyle et al, 2007; Cotgreave, 1996), it shows the value of the professional development focus of FSFE for staff.

What was most striking about the staff responses in this context was that—irrespective of prior teaching experience—all staff respondents reported that their professional participation in FSFE had improved their teaching skills. The great majority (90%) felt FSFE had specifically improved their capacity for reflection on their own teaching practice, and most felt this experience had improved their overall teaching, substantially in some cases. Many staff reported that, after participating in FSFE, they had been encouraged to do more teaching and seek out further professional development around their teaching.

Staff also commonly reported that FSFE benefited their research, by inspiring future research endeavours, and/or their career pathway, by providing development of both practical and 'soft' skills. Almost two thirds of teaching staff specifically reported an increase in their own technical confidence and their knowledge of ecology, with half of these reporting a significant increase (Fig. 5).

About two thirds of staff respondents also reported that participating in FSFE had resulted in a notable or significant increase in their professional networks. Staff identified a range of the longer-term benefits that had come from these broadened professional relationships, including unique opportunities to gain mentorship, initiate collaborations, and strengthen previously limited links with specific researchers.

##### Developing skills in reflective practice

Students were assessed on four elements of reflective writing. From the outset of the course students scored consistently high on ability to convey descriptive detail in their writing, which is appropriate as this is the most basic attribute of reflective writing (Moon, 1999). By contrast, between the mid- and end-of-course assessments there were statistically significant increases in scores for sophisticated and more effective reflective practice including emotive engagement with their experiences ($P=0.01$), ability to critically reflect, evaluate and analyse ($P<0.001$), and ability to reflect on the value of reflection (meta-reflection, $P<0.001$). Overall, and especially in students who attended both Intermediate and Advanced iterations, the journals demonstrated a clear shift from 'reflection *on* action' to 'reflection *in* action' which Schon (1983) considered the core of 'professional artistry', and Findlay (2008) described as indicating an expert who acts "both intuitively and creatively [as they] revise, modify and refine their expertise". This analysis of the reflective journal data was supported by the student survey findings: for example, most respondents (76%) reported a greater capacity to reflect on their own learning in their subsequent university courses after completing FSFE.

#### Literature cited

Bates D., Mächler M., Bolker, B., Walker, S. (2015). Fitting linear mixed-effects models using lme4. *Journal of Statistical Software*, 67(1), 1–48. doi: 10.18637/jss.v067.i01.

Beltran, R.S., Marnocha, E., Race, A., Croll, D.A., Dayton, G.H., & Zavaleta, E.S. (2020). Field courses narrow demographic achievement gaps in ecology and evolutionary biology, *Ecology and Evolution*, 10(12), 5184-5196. doi.org/10.1002/ece3.6300.

Boyle, A., Maguire, S., Martin, A., Milsom, C., Nash, R., Rawlinson, S., Turner, A., Wurthmann, S., & Conchie, S. (2007). Fieldwork is good: The student perception and the affective domain, *Journal of Geography in Higher Education*, 31:2, 299-317, doi: 10.1080/03098260601063628

Cotgreave, P. (1996). Fertile fields of study. *Times Higher Education*, May 17. https://www.timeshighereducation.com/news/fertile-fields-of-study/93654.article
Cotton, D.R. & Cotton, P (2009). Field biology experiences of undergraduate students: the impact of novelty space. *Journal of Biological Education*, *43*(4), 169-174. doi: 10.1080/00219266.2009.9656178

Moon, J. (1999). *Reflection in Learning and Professional Development: Theory and practice*. Abingdon, UK: Routledge-Falmer.

R Core Team (2021). R: A language and environment for statistical computing. R Foundation for Statistical Computing, Vienna, Austria. https://www.R-project.org/

Schon, D.A. (1983). The Reflective Practitioner. New York: Basic Books.

Finlay, L. (2008). Reflecting on 'reflective practice'. Practice-based Professional Learning Paper 52, The Open University.

Temponi, C. (2005), Continuous improvement framework: implications for academia, *Quality Assurance in Education*, 13(1), 17-36. doi:10.1108/09684880510578632
