## Supplementary File 3 for "An innovative approach to using an intensive field course to build scientific and professional skills"

### Nicotra et al.

#### Supplement 3: Survey Questions

These surveys were regulated by ANU Human Research Ethics Protocol 2015/553. Responses were anonymous and participants were notified that they could contact an independent researcher in confidence (see last question) if they desired.

**Evaluation (Students)**
1. In which course(s) did you participate?
2. When you FIRST attended the course, at which stage were you in your SCIENCE degree?
3. In which course were you enrolled?
4. In which degree program were you enrolled? (include both degrees if double)
5. Since you participated in the course(s), have you graduated?
6. What have you been doing since graduation? e.g. please tell us if you are pursuing a further degree (Honours, Master, PhD) or are employed (e.g. your job title) and if the role is in ecology, biology or another field
7. Field trips are both academic and social experiences. What are your clearest memories of the Field Studies in Functional Ecology course(s) you attended? Please feel free to include experiential memories (positive or negative) as well as those connected with academic or other learning.
8. How much did FSFE increase ...
   a) ... your INTEREST in ecology and ecosystems?
   b) ... your INTEREST in research?
   c) ... your KNOWLEDGE of ecology?
   d) ... your KNOWLEDGE of the ecosystems you visited?
   e) ... your TECHNICAL CONFIDENCE with using lab and field equipment in your studies?
   f) ... your SKILLS in formulating research questions?
   g) ... your SKILLS in planning research methodology?
   h) ... your CAPACITY to work effectively in groups?
   i) ... your CONCEPT of yourself as a SCIENTIST?
   j) ... your capacity to REFLECT on your learning in later courses?
   k) ... your NETWORKS of students and staff at the Research School of Biology?

9a. Did you contribute to writing a research paper in a published Studies in Functional Ecology journal?
9b. What did you feel about this opportunity to write and/or collaborate on a published paper?
10. After participating in the field trip, did you find any of the experiences continued to affect the way you studied? For example, in what ways did you find yourself more or less able to 'think like a scientist' or understand research?
11. Specifically, after the field trip, in what ways did you find yourself more or less able to work in groups?
12. In your opinion, what were the most useful activities/aspects of the course?
13. Please tell us if there were any aspects of the field trip(s) that you attended that you would have liked to have been different.
14. What impact do you think your participation in the field trip(s) had on your subsequent CHOICE of courses in your degree?
15. What impact do you think your participation in the field trip(s) had on your subsequent PERFORMANCE/ SUCCESS in courses in your degree?
16. Finally, is there anything else about your participation/experience in the course Field Studies in Functional Ecology that you would like to share? (If you would rather talk about these aspects confidentially, please feel free to contact [email])

**Evaluation (Staff)**

1a. Did you participate in any course(s) as an enrolled STUDENT?

1b. When you FIRST attended the course, at which stage were you in your SCIENCE degree?

1c. In which course(s) were you enrolled?

1d. In which degree program were you enrolled? (include both degrees if double)

1e. Since you participated in the course(s), have you graduated?

1f. What have you been doing since graduation? e.g. please tell us if you are pursuing a further degree (Honours, Master, PhD) or are employed (e.g. your job title) and if the role is in ecology, biology or another field

2. In which courses did you participate as a TUTOR, CO-CONVENOR and/or RESOURCE PERSON?

3. Which institution are you from?

4. What was your position in the course(s)?

5. Before your first Field Studies in Functional Ecology (FSFE) field course, how much experience did you have in supporting residential field courses/trips as TEACHING/RESOURCE staff?

6. Before your first FSFE field course, how much professional development experience did you have in TEACHING?

7. Field trips are both academic and social experiences. What are your clearest memories of the FSFE field trip(s) you attended? Please feel free to include experiential memories (positive or negative) as well as those connected with academic or other learning.

8. In your opinion, what were the most useful activities/aspects of the course for STUDENTS?

9. For the next set of questions, please bear in mind your own base rate of interest/knowledge when you first started TEACHING into the course. How much did your involvement in the FSFE course INCREASE ...

a) ... your INTEREST in research?
b) ... your KNOWLEDGE of ecology?
c) ... your KNOWLEDGE of the ecosystems you visited?
d) ... your TECHNICAL CONFIDENCE with using lab and field equipment?
e) ... your SKILLS in teaching?
f) ... your CAPACITY for effective teamwork?
g) ... your CONCEPT of yourself as a SCIENTIST?
h) ... your capacity to REFLECT on your teaching?
i) ... your NETWORKS of students and staff at the Research School of Biology and/or NTU?

10. Did you contribute to reviewing a research paper in a published Studies in Functional Ecology journal?

11. What did you feel about this opportunity for students to publish research?

12. Could you describe any ways in which your participation in the FSFE course(s) benefited you specifically in subsequently participate in any teaching professional development, or do you now do more teaching?

13. Could you describe any ways in which your participation in the FSFE course(s) benefited you specifically in terms of PROFESSIONAL RELATIONSHIPS with the other staff?

14. Could you describe any ways in which your participation in the FSFE course(s) benefited you specifically in terms of your own RESEARCH and/or CAREER path?

15. Please tell us if there were any aspects of FSFE that you would have liked to have been different.

16.Finally, is there anything else about your participation/experience in FSFE that you would like to share? (If you would rather talk about these aspects confidentially, please feel free to contact [email])
